# Brain transcriptome of gobies inhabiting natural CO2 seeps reveal acclimation strategies to long-term acidification

**DOI:** 10.1101/2022.09.18.508416

**Authors:** Sneha Suresh, Alice Mirasole, Timothy Ravasi, Salvatrice Vizzini, Celia Schunter

## Abstract

Ocean acidification (OA) is known to affect the physiology, survival, behaviour, and fitness of various fish species with repercussions at the population, community, and ecosystem levels. Some fish species, however, seem to acclimate rapidly to OA conditions and even thrive in acidified environments. The molecular mechanisms that enable species to successfully inhabit high CO_2_ environments has not been fully elucidated especially in wild fish populations. Here, we used the natural CO_2_ seep in Vulcano Island, Italy to study the effects of elevated CO_2_ exposure on the brain transcriptome of the anemone goby, a species with high population density in the CO_2_ seep and investigate their potential for acclimation. When compared to fish from environments with ambient CO_2_, gobies living in the CO_2_ seep showed differences in expression of transcripts involved in ion transport and pH homeostasis, cellular stress, immune response, circadian rhythm, and metabolism. We also found evidence of potential adaptive mechanisms to restore the functioning of GABAergic pathways, whose activity can be affected by exposure to elevated CO_2_ levels. Our findings indicate that gobies living in the CO_2_ seep may be capable of mitigating CO_2_ induced oxidative stress and maintaining physiological pH while meeting the consequent increased energetic costs. The conspicuous difference in expression of core circadian rhythm transcripts could provide an adaptive advantage by increasing flexibility of physiological processes in elevated CO_2_ conditions thereby facilitating acclimation. Our results show potential molecular processes of acclimation to elevated CO_2_ in gobies enabling them to thrive in the acidified waters of Vulcano Island.

## Introduction

Global change driven by anthropogenic activities is having profound effects on marine ecosystems with rising carbon dioxide (CO_2_) levels being one of the most crucial problems (Doney et al., 2009, 2012; Hofmann et al., 2010; Kroeker et al., 2010). Oceans are major carbon sinks and have absorbed nearly 30% of anthropogenic CO_2_ emissions since the 1980s resulting in ocean acidification (OA; Bindoff et al., 2019). Exposure to CO_2_ levels predicted to occur by the year 2100 (≥700 µatms) is known to impact the physiology, growth rate, development, survival, behaviour, and reproduction in various species of fish (Domenici et al., 2012; Esbaugh, 2017; Ferrari et al., 2012; Hamilton et al., 2017; Hamilton et al., 2014; Rachael M. Heuer & Grosell, 2014; Munday et al., 2013; Radford et al., 2021; Simpson et al., 2011). This can have far-reaching consequences on species interactions, population sustainability and ultimately impact ecosystem functioning (Fabry et al., 2008; Ferrari et al., 2011; Kroeker et al., 2010; Munday et al., 2010).

Most knowledge about these impacts of ocean acidification (OA) on teleosts, however, comes from short-term laboratory-based studies and mostly only account for short-term effects of OA. Accurate assessment of the future biological impacts of OA on species and populations requires consideration of their potential for acclimation and adaptive evolution (Sunday et al., 2014). The effects of elevated CO_2_ are also known to be variable both within and across species. Inter-individual variability in CO_2_ tolerance has been reported both in laboratory studies and in the wild (Cattano et al., 2018; Lehmann et al., 2022; Monroe et al., 2021; Munday et al., 2013; Paula et al., 2019; Welch & Munday, 2017). The underlying reason for the observed variable effect is not fully known but could be the result of specific genetic and epigenetic signatures or environmental factors or a combination of those. Environmental factors such as nutrient gradients, species composition, habitat modifications, and predation risk influence the effects of elevated CO_2_ on various aspects of fish physiology and behaviour (Ferrari et al., 2017; Mirasole et al., 2019; Nagelkerken et al., 2015; Zunino et al., 2019). Studies on wild fish populations that naturally inhabit high CO_2_ environments provide a unique opportunity for integrating the direct and indirect effects of elevated CO_2_ exposure on fish and investigate the molecular mechanisms enabling acclimation.

Volcanic CO_2_ seeps in the ocean, where emission of CO_2_-rich fluids locally reduce pH, have been used as natural analogues to study the effects of long-term exposure of fish to elevated CO_2_ in their natural environment. Shifts in habitat, trophic diversity and fish assemblages at CO_2_ seeps have been reported which could obscure some of the negative effects of OA on fish (Fabricius et al., 2011; Hall-Spencer et al., 2008; Mirasole et al., 2019; Nagelkerken et al., 2015; Vizzini et al., 2017). For example, predator recognition ability and risk behaviour of species living in CO_2_ seeps was found to be correlated with predator abundance at the seeps, with perception of predator threat to be unaffected by elevated CO_2_ in high predator risk environments (Cattano et al., 2017) and low-risk predation environments offsetting any negative impacts of CO_2_ induced impairment in predator recognition (Munday et al., 2014). For example, decrease in predator abundance at the CO_2_ seep could offset increased mortality of some fish species as a consequence of sensory and behavioural impairment resulting from exposure to elevated CO_2_ levels (Munday et al., 2014). Habitat shifts is another factor that can work antagonistically with CO_2_ to influence species behaviour. Gobies from the CO_2_ seeps in Vulcano Island, Italy living in more open habitats showed reduced predator escape behaviour, however, the same was not observed in gobies living in more sheltered sites indicating the influence of habitat on species response to elevated CO_2_ exposure (Nagelkerken et al., 2015). Similarly, habitat shifts resulting in net resource enrichment at CO_2_ seeps seems to benefit certain fish species by providing sufficient food to meet the increased energy demands of living in a high CO_2_ environment (Connell et al., 2013; Mirasole et al., 2019; Nagelkerken et al., 2015). It is therefore important to consider the indirect effects of OA while making inferences about the adaptive potential of species to future OA conditions and volcanic CO_2_ seeps provide the perfect opportunity to do so.

Prolonged exposure to elevated CO_2_ at the seeps may enable fish to acclimate through physiological and behavioural compensation resulting in long-term survival and subsequent adaptation. Indeed studies using transcriptomics to determine gene expression patterns in the brain and gonads of fish from natural CO_2_ seeps have revealed plasticity and adaptive selection to elevated CO_2_ (Kang et al., 2022; Petit-Marty et al., 2021). These findings suggest that species that are continuously exposed to elevated CO_2_ levels can develop mechanisms to mitigate negative effects of CO_2_ exposure and have the potential to adapt to acidified environments. However, given the relatively small spatial range of CO_2_ seeps compared with the pelagic larval dispersal distance of most marine species, the population in the CO_2_ seep will mostly come from a heterogeneous larval pool. The recently settled larvae in the seep must therefore rapidly respond to the acidified environment via phenotypic plasticity, resulting in physiological adjustments to ensure survival. These plastic responses established during development could be crucial to allow organisms to acclimate and eventually lead to adaption to environments with elevated CO_2_ levels on a local scale (Munday, et al., 2013; Sunday et al., 2014). As the CO_2_ levels continue to rise developmental plasticity could be a crucial process facilitating the establishment of adaptive processes across species enabling them to survive in the increasingly acidifying oceans of the future.

While several studies have investigated the effects of elevated CO_2_ on fish physiology, behaviour, and molecular processes, we are only starting to learn about the effects of CO_2_ on brain function in wild fish species. Here, we sequenced and analysed the brain transcriptome of the anemone goby, *Gobius incognitus* to identify gene expression patterns underlying acclimation to elevated CO_2_ in the wild population. This benthic fish species has a very limited home range making it an ideal model to study the effects of elevated CO_2_ exposure *in situ*. Previous studies have reported potentially negative physiological effects such as changes in otolith morphology and skeletal composition in *Gobius* from the natural CO_2_ seeps at Vulcano Island (Mirasole et al., 2017, 2021), however, they still maintain higher population densities at the seeps (Nagelkerken et al., 2015). This could mean that gobies have developed alternative strategies/mechanisms to offset the negative effects of elevated CO_2_ exposure. In fact, swimming activity and predator perception of *Gobius incognitus* living in natural CO_2_ seeps in Vulcano Island was not affected by elevated CO_2_ indicating higher tolerance of this species to changing ocean conditions (Spatafora et al., 2022). By examining brain gene expression patterns of the anemone goby from the natural CO_2_ seeps in Vulcano Island, Italy we aim to identify molecular signatures of acclimation that enable this species to mitigate negative effects of elevated CO_2_ and thrive in the acidified waters of Vulcano Island.

## Methods

### Study area and sample collection

*Gobius incognitus* samples were collected by SCUBA diving on 30^th^ September 2018 at a depth of 1-2 meters with small hand nets from two locations in Levante Bay, Vulcano Island (Italy): a low pH site (38.419 °N, 14.961 °E; pH 7.80 ± 0.09) and a control site (38.420 °N, 14.965 °E; pH 8.19 ± 0.03) located around 600 m from the low pH site (Figure 1). The control site was similar to the low pH site in terms of orientation (south-east), depth and habitat (sandy with beds of *Cymodocea nodosa* seagrass). A total of eight individuals were sampled at each site at the same time of day (between 11:00 am and 12:00 pm). The total length of all fish was measured in the field, immediately after which they were euthanized on ice and the brain tissues were dissected, preserved in RNAlater, and stored at -80°C until further processing. The age of all sixteen individuals was estimated using the Von Bertalanffy growth function (VBGF; von Bertalanffy., 1938) which is based on a bioenergetic expression of fish growth. Size-at-age data can be used to fit parameters of a growth function, such as the VBGF. In our case, we extrapolated the age of our analysed Gobius specimens using the growth data from the work of Sasal et al., 1996. On average all the sampled fish were four cm long and two years old (Supplementary Table S1).

**Figure 1:**
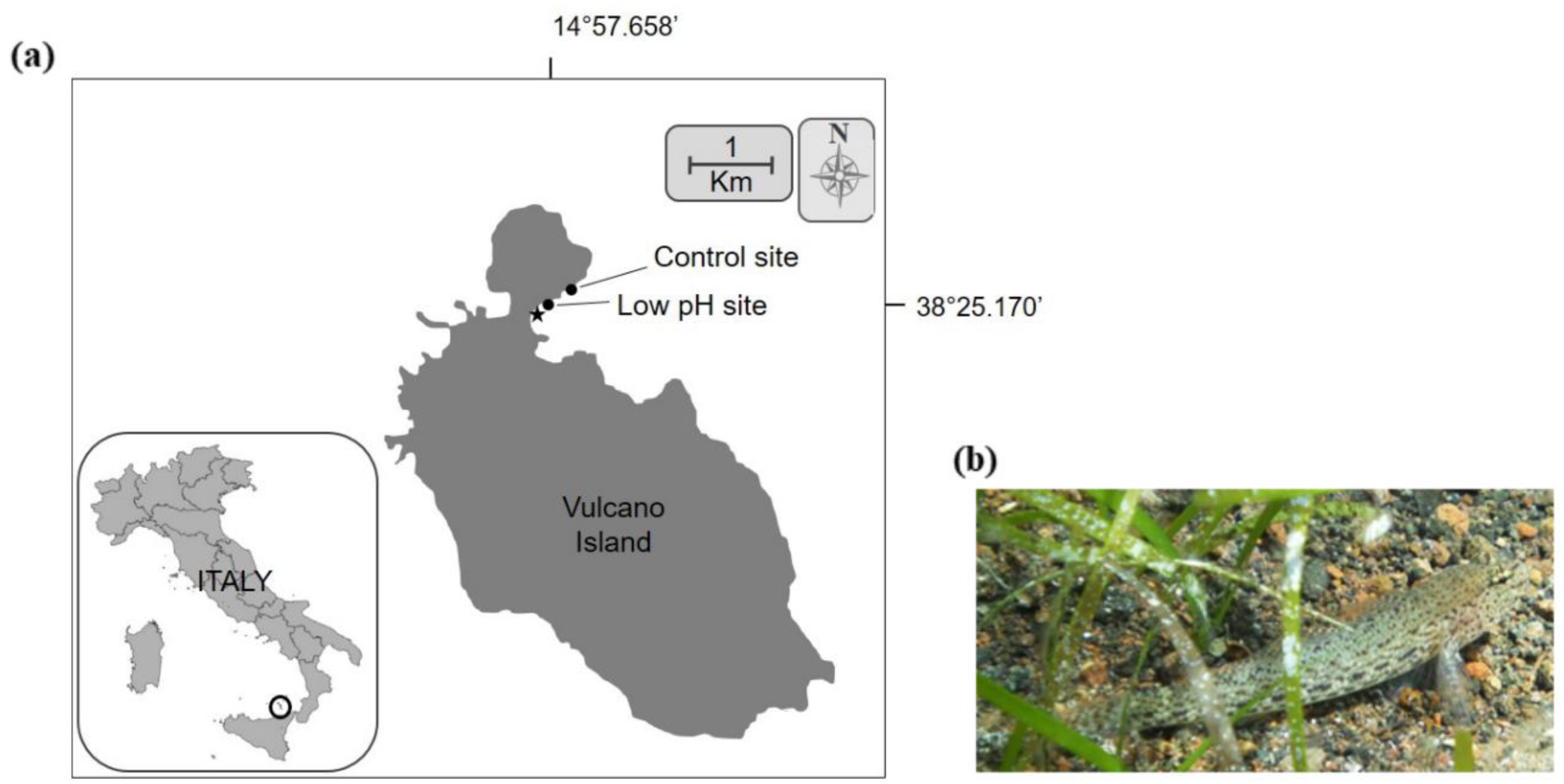
(a) Map showing the sampling sites in Levante Bay, Vulcano Island. The star indicates the location of the primary vent and the circles indicate the low pH site and the control (ctrl) site. (b) Picture of the study species *Gobius incognitus* in the wild.

### RNA extraction and sequencing

Frozen whole brains were homogenized in RLT-lysis buffer (Qiagen) for 30s at 30Hz/s using Qiagen TissueLyzer II with 4-5 single-use silicon beads per sample and total RNA was extracted using the Qiagen RNeasy Micro Kit. On-column DNase digestion was performed using RNase-Free DNase1 (Qiagen) following the manufacturer’s instructions, to remove any DNA contamination. RNA concentration and quality was determined using Qubit and Agilent 2100 Bioanalyzer respectively (mean RIN = 7.79) following which RNA-Seq libraries were built using the KAPA Stranded mRNA-Seq Kit with 100 ng total RNA as input from each of the 16 individuals. The samples were subsequently sequenced on an Illumina NovaSeq 6000 to get 151 bp, paired-end reads at the Centre for PanorOmics Sciences, The University of Hong Kong. An average of 33 ± 2.8 million read pairs were obtained from the 16 RNA libraries sequenced (Supplementary Table S2).

### Sequence processing and primary de novo transcriptome assembly

The quality of all the sequencing reads was determined using FastQC v0.11.5 (Andrews, 2010). To ensure high quality of all reads, we trimmed adapters and sequences with an average Phred-scaled quality score below 30 using fastp v0.20.0 (Chen et al., 2018) with a 4-base sliding window approach. Sequences less than 40 bp in length after trimming were removed. The trimmed sequences were further filtered to remove potential contaminants, ambiguous bases, and rRNA sequences. Potential contaminant sequences were removed using kraken v2.0.8-beta (Wood & Salzberg, 2014) with a confidence score of 0.3 using the bacterial, fungal, and viral databases from NCBI RefSeq as reference. For sequences containing ambiguous base calls (Ns), the longest sub-sequence of at least 40 bp without any ambiguous bases was extracted and retained if it was at least half the original sequence length and had an average Phred-scaled quality score above 30 using the ‘filter_illumina’ script from DRAP (Cabau et al., 2017). Reads originating from ribosomal RNA (rRNA) were identified and removed by mapping the sequences to the SSUParc (SILVA_138_SSUParc_tax_silva.full_metadata.gz) and LSUParc datasets (SILVA_132_LSUParc.full_metadata.gz) from SILVA (Quast et al., 2013) using Bowtie2 v2.4.1 (Langmead & Salzberg, 2012) in the very sensitive, local mode. An average of 26 ± 2.5 million adapter free, filtered, quality-trimmed, and decontaminated paired reads were finally retained after the stringent filtering process from all 16 samples (Supplementary Table S2). These high quality reads were subsequently merged and assembled *de novo* using Trinity v2.8.4 (Grabherr et al., 2013) within DRAP v1.92 (parameters: -c ’-x’; -- no-trim; -m bwa; --no-rate) using the pooled-assembly strategy resulting in a primary transcriptome assembly consisting of 633,517 contigs.

### *De novo* transcriptome assembly optimization

Candidate coding regions (open reading frames (ORFs)) within the transcript contigs assembled by DRAP were identified using TransDecoder v5.5.0 (Haas et al., 2013) and only contigs with ORFs of at least 100 amino acids long were retained to prevent retention of false positive ORFs as done in previous studies on CO_2_ seeps (Kang et al., 2022; Petit-Marty et al., 2021). These contigs were then blasted against the UniProtKB/Swiss-Prot Release 2021_01 database (Apweiler, 2004) using the BLASTX algorithm (Blast+ v2.6.0) (Altschul et al., 1990) with an e-value cut-off 1e-05 and retaining only one blast hit per contig (--max_target_seqs = 1). For each assembled contig only the best ORF was retained prioritized first based on blast homology and then based on ORF length (--retain_blastp_hits; --single_best_only) after which 113,895 contigs were retained. The quality of the selected contigs was further assessed using the runAssesment module within DRAP v1.92 and 68 putative chimeric contigs were removed. The high-quality RNA-seq reads were then re-mapped to the filtered transcriptome assembly using Bowtie2 v2.4.1 in the sensitive mode and disallowing any discordant and unpaired alignments, but without any upper limit on the number of alignments. To identify and cluster individually assembled contigs belonging to the same gene, the alignment results were used by Corset v1.09 (Davidson & Oshlack, 2014) to hierarchically cluster contigs based on the proportion of multi-mapped reads and the expression patterns. For each of the Corset generated clusters the longest transcript sequence was extracted using the fetchClusterSeqs.py script from Corset-tools (https://github.com/Adamtaranto/Corset-tools) and those clusters with no aligned reads were removed from the *de novo* transcriptome assembly resulting in 44,614 high quality contigs retained.

To further validate the transcriptome assembly, blast search (TBLASTX, e-value = 1e- 05, max_target_seqs = 1) was performed against the complete cDNA sequence of *Neogobius melanostomus* (round goby) obtained from Ensembl (release-104) (Howe et al., 2021), the phylogenetically closest species with a sequenced genome to *Gobius incognitus* (Thacker & Roje, 2011). To further reduce transcript redundancy and identify potential alternative transcripts that failed to cluster in the previous steps, the blast results were filtered to retain only the longest transcript for each blast hit to get the final *de novo* transcriptome assembly. Transcripts with no blast hits to *Neogobius melanostomus* were also retained in the final assembly as these could potentially be transcripts that are specific to our target species, *Gobius incognitus*. The final transcriptome assembly consisted of 43,349 non-redundant transcripts. The basic statistics of the transcriptome assembly determined using rnaQUAST v2.1.0 (Bushmanova et al., 2016), ‘stats.sh’ script from BBMap v38.87 (Bushnell, 2014) and ‘contig_ExN50_statistic.pl’ from Trinity accessory scripts (https://github.com/trinityrnaseq/trinityrnaseq/tree/master/util/misc), are shown in Supplementary Table S3 and Supplementary Figure S1.

### Transcriptome annotation and evaluation of assembly completeness

The *de novo* assembled transcriptome was compared against the UniProtKB/Swiss-Prot Release 2021_01 database using BLASTX (e-value = 1e-05 and HSP length cut-off = 33) resulting in 43.6% (24,454) of the assembled transcripts with blast hits to known proteins. Gene Ontology (GO) mapping and annotation was then carried out within OmicsBox v1.4.11 (Gö Tz et al., 2008). InterProScan was used to identify the protein domains within the transcript sequences and the InterProScan GOs were merged with the GO annotations resulting in a total of 24,875 (42.6%) annotated transcripts. For contigs that did not blast to the Swiss-Prot database, the Ensembl gene ID obtained by blasting against *Neogobius melanostomus* was retained. Combining the annotation results from OmicsBox and the additional 9,362 transcripts with blast hits to the cDNA sequence of *Neogobius melanostomus*, 78.9% (34,237) of the transcripts in the final *de novo* assembly has a functional annotation (Supplementary Table S3 and S4).

To incorporate the transcript expression levels in evaluating the contiguity of the *de novo* assembly, the ExN50 (expression-informed N50) metric was determined. ExN50 (expression-informed N50) represents the N50 value for the most highly expressed transcripts representing x% of the total normalized expression data and is more appropriate for transcriptome assemblies than the conventional N50 statistic (Galachyants et al., 2019). A peak of the ExN50 value at higher x percentiles indicates the effectiveness of the de novo assembly pipeline to assemble longer transcripts (Sahraeian et al., 2017). The expression levels of the transcripts were quantified using RSEM v1.3.1 (Li & Dewey, 2014) and the ExN50 value was calculated across different expression percentiles using the TMM normalized expression values. The *Gobius incognitus reference* transcriptome assembled here has a sufficiently high coverage of the longer transcripts indicated by a peak of ExN50 value at E86 (Supplementary Table S5, Supplementary Figure S2). The level of completeness of the transcriptome assembly was determined using BUSCO v4.1.3 (Manni et al., 2021) using the Actinopterygii_odb10 and Eukaryota_odb10 sets of Benchmarking Universal Single-Copy Orthologs (BUSCO) which revealed high representation of both the Actinopterygian core genes and the Eukaryota core genes. Specifically, 76.4% and 91.4% of Actinopterygii core genes and Eukaryota core genes respectively, were recovered completely. An additional 7.7% of Actinopterygii core genes and 7.1% of Eukaryota core genes were recovered partially (Supplementary Figure S3) indicating sufficiently good assembly and annotation completeness, similar to other *de novo* assemblies (Pomianowski et al., 2021; Wong et al., 2019).

### Differential expression analysis

The high-quality, filtered RNA-seq reads were re-mapped to the final transcriptome assembly using Bowtie2 v2.4.1 in the sensitive mode, disallowing any discordant and unpaired alignments and using additional parameters (--dpad 0 --gbar 99999999 --mp 1,1 --np 1 --score-min L,0,-0.1) to ensure the resulting alignment files are compatible with RSEM, which was used to quantify the transcript expression levels. On average 72% of the RNA-Seq reads mapped back to the assembled transcriptome (Supplementary Table S6) indicating sufficiently high quality of the *de novo* transcriptome. Principal component analysis (PCA) was performed to determine the overall transcript expression patterns and identify any outlier samples in R v4.1.1 (R Core Team, 2021) following which differential expression analysis was done using DESeq2 v1.32.0 (Love et al., 2014). The transcripts were considered to be significantly differentially expressed between the control and CO_2_ seep sites if the FDR corrected p-value (padj) was less than 0.05. Additionally, to remove further false positives, we set other filters to absolute log2 fold change (abs log2FC) greater than 0.3 and baseMean greater than 10 as done in previous studies on brain fish transcriptomics (Schunter et al., 2016, 2018). Enrichment of GO functional categories among the significant DE genes were determined using Fisher’s Exact Test in OmicsBox v1.4.11 (*OmicsBox – Bioinformatics Made Easy, BioBam Bioinformatics, March 3*, 2019) with an FDR threshold of 0.05, using the option “reduce to most specific” to reduce the redundancy in the enriched GO categories identified. The heatmap showing relative transcript expression patterns across samples was created using the pheatmap package v1.0.12 (where transcripts with similar expression levels were clustered together) and all other figures were created using the ggplot2 package v3.3.6 in R v4.2.1.

### SNP calling and detection of outlier loci

SNP calling and outlier detection was performed to determine if there were any genetic differences between the samples from the CO_2_ seep and control sites. The previously obtained BAM files were marked for duplicate reads using MarkDuplicates (Picard) and reads containing Ns in their cigar string were split using SplitNCigarReads within the Genome Analysis Toolkit (GATK v4.3.0.0) (Depristo et al., 2011). SNPs within the samples were called using HaplotypeCaller within GATK and filtered using the VariantFiltration tool (parameters: -window 35 -cluster 3 --filter-name FS -filter “FS>30.0” --filter-name QD -filter “QD<2.0”). The resulting VCF files from all 16 individuals were merged and further filtered to remove all indels and retain only SNPs with a Phred- scaled quality score of 30 and minor allele frequency (MAF) ≥ 0.2. Additionally, the minimum and maximum depth cut-off was set to 10 and 70 respectively per site and per genotype to account for possible mapping/assembly errors and all sites where over 25% of the individuals are missing a genotype were removed. Discriminant Analysis of Principal Components (DAPC; Jombart et al., 2010) was used to determine genetic structuring between the samples and to evaluate if particular variants were potentially under selection, outlier loci detection was performed in BayeScan v2.0 software (Foll & Gaggiotti, 2008) with the assumption that the individuals from the control and CO_2_ seep sites belong to two different populations.

## Results

### Brain transcriptome analysis

A total of 43,301 transcripts were expressed across all 16 samples with fish from the CO_2_ seep site showing more variability in transcript expression among individuals compared to fish from the control site (Figure 2). There was no significant difference in the total body length (Welch’s t-test, t(19.48) = 0.43, p = 0.67) and age (Welch’s t-test, t(19.64) = 0.47, p = 0.64) of the fish sampled from the low pH site and control site. Therefore none of the measured biological factors drive the observed transcript expression patterns. When comparing transcript expression levels between the control and CO_2_ seep sites, 996 transcripts were identified to be significantly differentially expressed (DE) (padj < 0.05, abs log2FC > 0.3, baseMean > 10; Supplementary Table S7). Majority of these (68%) were expressed at higher levels in fish from the CO_2_ seep site compared to the control site (Supplementary Figure S4).

**Figure 2:**
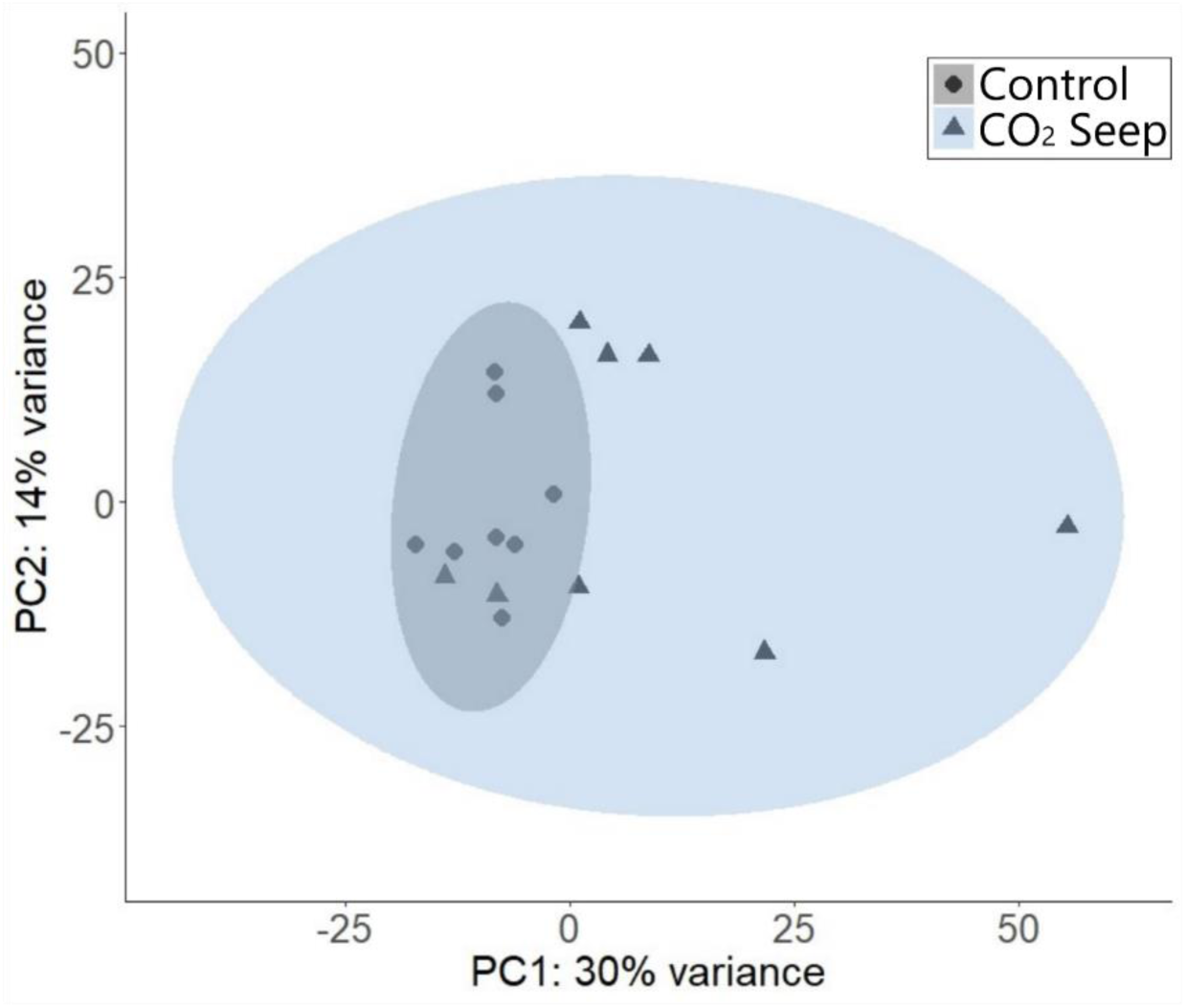
Overall transcript expression pattern across the 16 samples from the natural CO_2_ seep and control site at Vulcano Island. Principal component analysis (PCA) was performed using the regularized log transformed (rlog) counts of all expressed transcripts. The circles represent the individuals from the control site and the triangles represent the individuals from the low pH site. The blue and grey region is the 95% confidence interval of the samples from the CO_2_ seep and control sites respectively.

### Functional analysis of DE transcripts

Functional enrichment analysis identified 434 GO terms to be enriched among the DE transcripts (Supplementary Table S8). The transcripts associated with each enriched GO term was further categorised into six broader functional groups (acid-base and ion homeostasis, neural activity and related effects, circadian rhythm, metabolism, cellular stress response, and immune response) based on their functional description from the NCBI gene database (https://www.ncbi.nlm.nih.gov/gene/) and UniProt knowledgebase (UniProtKB; https://www.uniprot.org/) (Figure 3, Supplementary Table S9).

**Figure 3:**
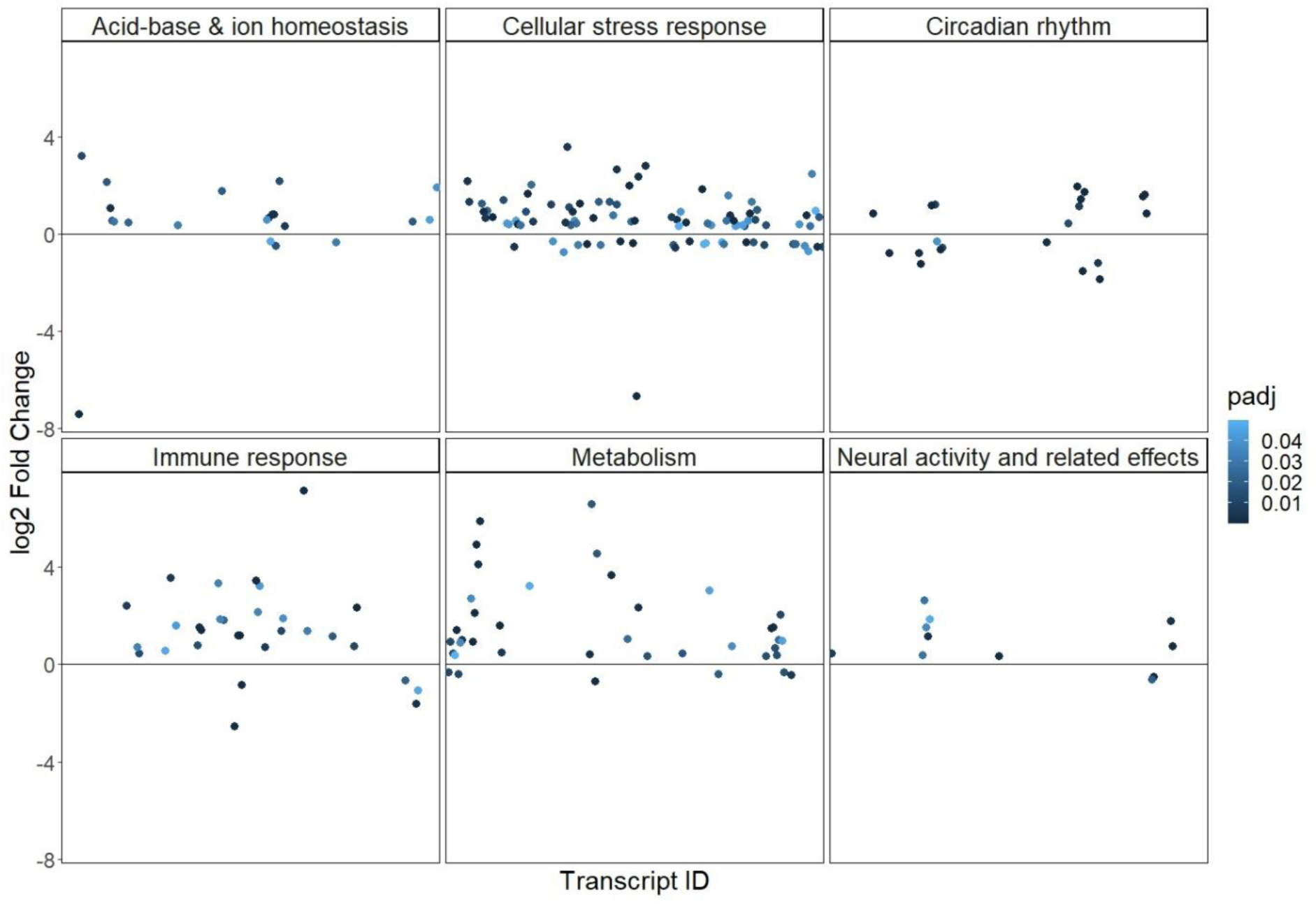
Functional groups associated with the differentially expressed transcripts. Each circle represents one transcript, and the colour represents the FDR corrected p-value (padj) from the differential expression analysis.

The five most significant DE transcripts (with the lowest padj values) were all associated with the circadian rhythm. These include two core circadian genes Period circadian protein homolog 3 (*PER3*) and Orphan nuclear receptor RVR (*NR1D2*), two nuclear hormone receptor genes involved in transcriptional regulation of core circadian genes Nuclear receptor ROR-alpha A (*RORA*) and Nuclear receptor ROR-beta (*RORB*), and a transcriptional regulator known to regulate circadian rhythm genes Nuclear factor interleukin-3-regulated protein (*NFIL3*) (Supplementary Figure S4). In fact, all core clock transcripts were DE between the control and CO_2_ seep site; specifically, there was an overall downregulation of repressors of the circadian rhythm and upregulation of the activators (Figure 4). Furthermore, *NOCT* a circadian deadenylase that functions as a post transcriptional modulator in circadian control of metabolic genes was upregulated in individuals from the CO_2_ seep (Supplementary Table S9).

**Figure 4:**
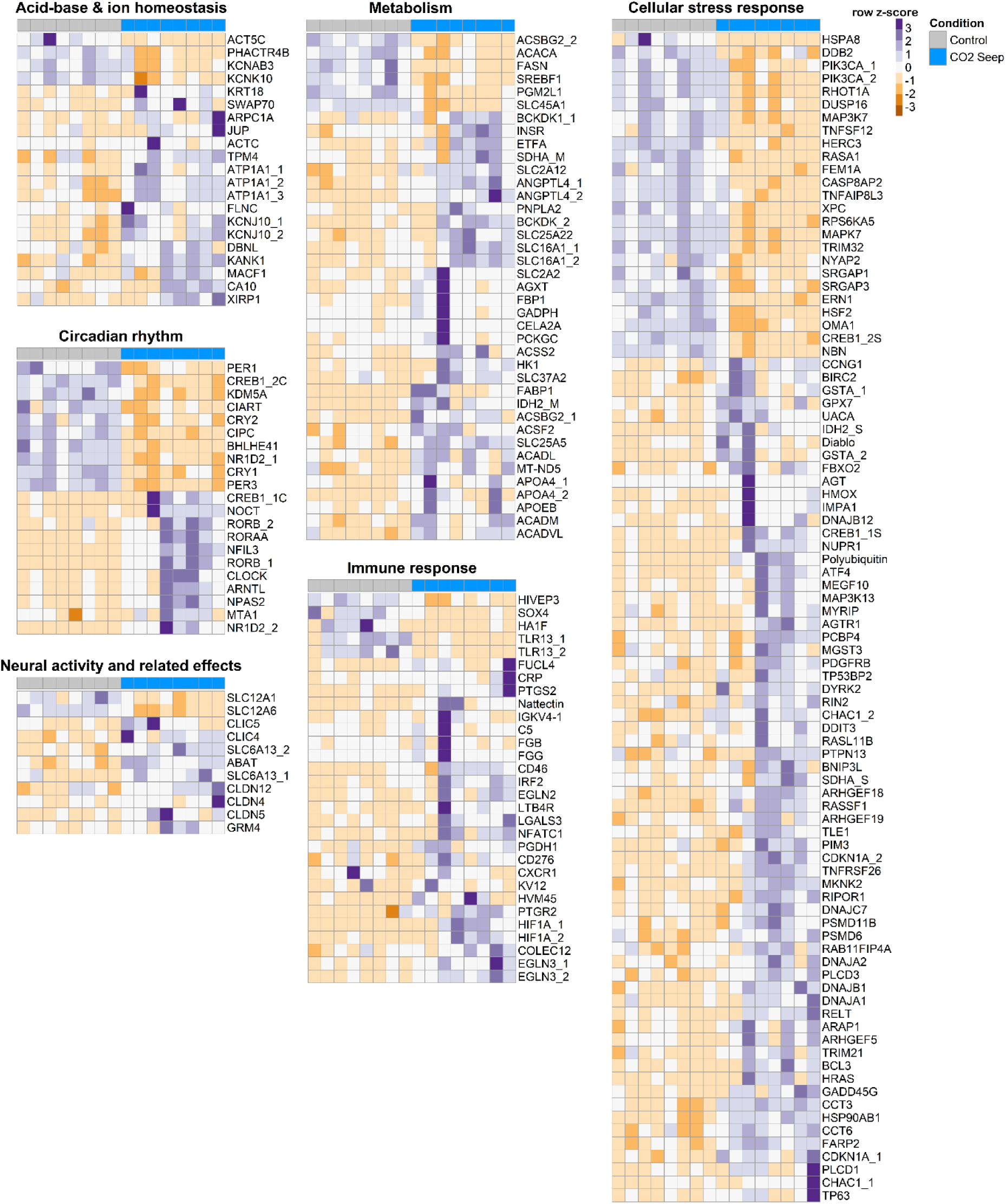
Relative expression levels of transcripts with enriched functions of fish from control and CO_2_ seep sites. Each row represents a transcript, and the row z-score is calculated from the log2 transformed transcripts per million (TPM) values for each transcript. The transcripts are clustered based on expression levels.

Various transcripts involved in cellular stress response was also DE in our study. In fact this group had the highest number of DE transcripts majority of which were upregulated in individuals from the CO_2_ seep (Figure 4). This included transcripts known to play a role in the production of reactive oxygen species (ROS), transcripts involved in phosphatidylinositol 3- kinase (PI3K) activity and signalling, DNA damage repair, unfolded protein response and ubiquitination, molecular chaperones including heat shock proteins (HSPs), cell cycle control, apoptosis, various members of the Ras superfamily of small GTPases and transcripts associated with redox-sensitive signalling pathways such as the NF-*k*B and MAPK signalling. Notably, there was also an overall increase in the expression of transcripts associated with cellular detoxification and redox regulation (such as glutathione transferase, glutathione peroxidase, heme oxygenase, isocitrate dehydrogenase, and succinate dehydrogenase), cellular response to hypoxia, immune response and immune system function in individuals from the CO_2_ seep site. Of particular interest was the *galactose-specific lectin nattectin*, a C-type lectin which is involved in innate immune responses as it had the highest fold-change in expression among all DE transcripts (Supplementary Table S9).

Ion channels and transporters were another major functional group that was differentially regulated between fish from the CO_2_ seep and control sites. Notably, transcripts encoding the Na^+^/K^+^ transporting ATPase (*ATP1A1*) and carbonic anhydrase 10 (*CA10*), which play critical roles in ion-regulation and maintaining pH homeostasis, were upregulated in fish from the CO_2_ seep site (Figure 3). Additionally, various transcripts involved in actin cytoskeleton organisation were also DE (Supplementary Table S9, Figure 4). This could indirectly affect membrane transport as the cytoskeleton closely interacts with the plasma membrane. Changes in ionic composition across neuronal membranes can affect the activity of neurotransmitters such as GABA and glutamate and we found transcripts involved in degradation of GABA (*SLC6A13* and *GABAT*) and a transcript encoding the metabotropic glutamate receptor 4 (*GRM4*) to be differentially expressed in our study (Supplementary Table S9, Figure 4). Additionally, transcripts encoding chloride transporters and claudins, whose expression is affected by neural activity, were also DE (Supplementary Table S9, Figure 4).

Several transcripts involved in metabolism were also found to be differentially regulated between fish from control and CO_2_ seep sites. Transcripts involved in key metabolic processes that result in increased ATP production such as glycolysis, gluconeogenesis, TCA cycle, mitochondrial beta-oxidation, mitochondrial respiratory chain, lipid and amino acid metabolism were expressed at higher levels in gobies from the CO_2_ seep site. Interestingly, transcripts involved in fatty acid synthesis, an ATP consuming process were downregulated in individuals from the CO_2_ seep (Supplementary Table S9, Figure 4).

### Outlier loci detection

We evaluated the genetic background of the fish living in CO_2_ seep and control site. A total of 8,629 high-quality SNPs were obtained across all 16 individuals from the CO_2_ seep and control site. There was no genetic structuring found among the analyzed samples and no outlier was detected (Supplementary Figure S5). Therefore there was no evidence of genetic divergence between the individuals from the control and CO_2_ seep site. Gobies have a pelagic larval phase and hence the individuals from the CO_2_ seep and control sites come from a heterogeneous pool of larvae and there is no evidence of post-settlement processes between the sites.

## Discussion

Natural CO_2_ seeps provide an excellent opportunity to study the effects of long-term exposure of fish to elevated CO_2_ in their natural environment and determine their adaptive potential while also accounting for various indirect effects of ocean acidification (OA) such as habitat shifts and changes in species assemblages. Here, we investigated changes in the brain transcriptome of the anemone goby from a natural CO_2_ seep in Vulcano Island to identify molecular processes enabling acclimation to elevated CO_2_ in the wild. Previous studies at Vulcano Island have found limited behavioural effects of CO_2_ on this fish species and increased population size at the seep suggesting that this species has higher tolerance to elevated CO_2_ conditions possibly through differential regulation of gene expression enabling physiological and behavioural adjustments (Nagelkerken et al., 2015; Spatafora et al., 2022).

We found large differences in overall brain transcript expression levels between individuals from CO_2_ seep and ambient control sites. The expression patterns were more variable among individuals from the CO_2_ seep compared to those from the control site which is not due to the age or length of the fish. Additionally, the SNP analysis did not reveal any post-settlement selection processes which could be driving the increased variability of transcript expression levels in fish from the CO_2_ seep compared to those from the control site. Several previous studies have reported variability in species response to elevated CO_2_ (Cattano et al., 2018; Monroe et al., 2021; Munday et al., 2013; Paula et al., 2019; Welch & Munday, 2017), which could be due to variation in acclimatization strategies or epigenetic differences. The variation within this population could also result from inter-individual differences in capacity for plasticity when faced with environmental change (Forsman, 2014). Given that developmental plasticity is potentially the primary mechanism driving local adaptation within the CO_2_ seep, it is highly likely that the observed variation in transcript expression is resulting from post-settlement plastic responses. Further research is however needed to fully understand the underlying causes of individual variation in CO_2_ tolerance especially since intraspecific variability can be heritable (Welch & Munday, 2017) and hence is a key factor in influencing adaptive capacity of populations to future global change (Lehmann et al., 2022; Munday et al., 2013).

The majority of differentially expressed (DE) transcripts exhibited elevated expression in fish from the CO_2_ seep similar to what has been observed in gonads of the common triplefin from the CO_2_ seep in New Zealand, another species known to have increased population size in acidified environments (Petit-Marty et al., 2021). The gobies as well as the common triplefin seem to be able to elicit acclimation responses to elevated CO_2_ by differentially regulating transcripts associated with crucial functions which potentially enables them to maintain cellular homeostasis and thrive in acidified waters at CO_2_ seeps showing a more general adaptive capacity of species that thrive in these environments. Specifically, for the gobies 2.3% of the brain transcriptome involved in key functions such as acid-base and ion homeostasis, neurological function, circadian rhythm, metabolism, cellular stress response, and immune response were differentially expressed (DE) between fish from the CO_2_ seep and control sites.

Cellular Stress Response was the most gene rich response in gobies at the CO_2_ seep. We observed DE of various transcripts associated with redox sensitive pathways such as NF- *k*B and MAPK signalling as well as transcripts known to induce an increase in the intracellular concentration of reactive oxygen species (ROS). Therefore, gobies inhabiting the CO_2_ seep could experience changes in cellular redox balance resulting in oxidative stress (Poli et al., 2012; Zhang et al., 2016). They however also seem to be capable of counteracting stress-induced damage by upregulating transcripts involved in cellular detoxification and redox regulation. Oxidative stress in fish can trigger cell damage and apoptosis (Pizzino et al., 2017; Strader et al., 2020) and also cause conformational changes and oxidative damage to all macromolecules including DNA, proteins, and lipids thereby affecting transmission of genetic information and resulting in intracellular accumulation of unfolded and misfolded proteins (Juan et al., 2021) which is cytotoxic (Hetz, 2012). Differential regulation of transcripts involved in DNA damage response and DNA repair, unfolded protein response (UPR), chaperones such as the heat shock protein (HSP) family observed here suggest that gobies are capable of minimising oxidative damage on macromolecules caused by ROS. High constitutive expression of several universally conserved genes in the cellular stress response machinery in fish from the CO_2_ seep indicate an adaptive response to increase cellular resistance to oxidative stress induced cell damage.

Transcripts involved in innate and adaptive immune response and the complement system were found to be up-regulated in gobies from the CO_2_ seep site. Elevated levels of cellular stress can result in increased infections from opportunistic pathogens and hence activation of immune responses could be a protective mechanism to prevent potential infections in the face of increased physiological stress in environments with elevated CO_2_ levels (de Souza et al., 2016). Immune response genes are commonly differentially regulated in fish exposed to elevated CO_2_ (de Souza et al., 2016; De Souza et al., 2014; Machado et al., 2020) revelaing it as an important function that is regulated under elevated CO_2_ conditions. Additionally, two hypoxia-inducible transcripts, *HIF1A* and *EGLN2* were upregulated in individuals from the CO_2_ seep. While the primary function of these transcripts is to regulate cellular response to hypoxic conditions, their expression could also be enhanced in response to cellular stress. Increased mRNA expression of *HIF1A* in response to hypercapnia has been previously reported under stable oxygen levels (Machado et al., 2020). These results indicate that gobies living in the naturally acidified waters of Vulcano Island are able to elicit physiological compensatory mechanisms to counter the cellular stress induced by prolonged exposure to elevated CO_2_ levels.

Living in CO_2_ seeps also induced expression differences in various transcripts associated with ion transport and conserving pH balance. In particular, carbonic anhydrase (CA), an enzyme which plays a key role in CO_2_ excretion, acid-base balance, and ion- regulation was upregulated in gobies from the CO_2_ seep, which could facilitate increased CO_2_ excretion thereby preventing acidosis and also enhance transport of O_2_ to the brain (Rummer & Brauner, 2011). Additionally, transcripts encoding potassium channel transporters (*KCNAB3*, *KCNK10* and *KCNJ10*), and a Na^+^/K^+^ transporting ATPase (*ATP1A1*) were differentially expressed (DE) in the gobies. Acid-base balance and ion homeostasis in fish are closely linked processes as transport of H^+^ and HCO_3_^-^ ions are coupled to K^+^, Na^+^ and Cl^-^ transport (Gilmour & Perry, 2009). Regulation of ion transport is therefore key to ensure proper cellular functioning, maintain pH homeostasis and prevent respiratory acidosis which is a common effect of elevated CO_2_ exposure in fish (Heuer & Grosell, 2014; Ishimatsu et al., 2005). DE of transcripts encoding potassium channel proteins in our study could potentially be involved in enabling gobies living in the CO_2_ seep to sense ambient CO_2_ levels and in turn trigger downstream homeostatic responses to restore internal acid-base status as potassium channels are involved in CO_2_ chemoreception and sensing acid-base disturbance (Qin et al., 2010). Additionally, the increased expression of *ATP1A1* in fish from the CO_2_ seep may play an important role in pH homeostasis as the sodium and chloride gradient created by this enzyme affects the activity of other key secondary transporters involved in pH regulation (Casey et al., 2010; Singh et al., 2015; Tresguerres et al., 2020). Transcripts involved in actin cytoskeleton organisation was also DE which combined with DE of Ras GTPases, which are involved in remodelling the cytoskeleton, could imply changes in membrane-cytoskeleton interactions in individuals from the CO_2_ seep, in turn affecting membrane transport (de Curtis & Meldolesi, 2012). Therefore, gobies inhabiting the naturally acidified waters of Vulcano Island can efficiently regulate various ion transport mechanisms to buffer any intracellular pH changes caused by elevated environmental CO_2_ and thereby maintain cellular homeostasis.

We found subtle evidence of regulation of GABAergic signalling in gobies from the CO_2_ seep which could be a consequence of changes in ion channel activity as neural signalling pathways are reliant on ion transport and stable ion gradients. Two transcripts, *SLC6A13* and *GABAT* regulating the turnover rate of extracellular γ-aminobutyric acid (GABA), which is a major neurotransmitter in the brain, were upregulated in gobies from the seep. Given that GABA is not prone to enzymatic degradation, increased expression of these transcripts could be an adaptive response to regulate GABAergic signalling which is known to be altered in high CO_2_ environments as a result of pH buffering processes (Heuer et al., 2016; Hübner & Holthoff, 2013; Kang et al., 2022; Nilsson et al., 2012; Schunter et al., 2019). Furthermore, upregulation of the glutamate metabotropic receptor 4 (*GRM4*) in fish from the CO_2_ seep could be a compensatory response to reduce presynaptic GABA release (Bobeck et al., 2014; Flor et al., 1995). Additionally, transcripts involved in the transport of chloride ions, whose concentration gradient influences GABA activity, were DE, which could potentially regulate GABAergic neurotransmission (Jentsch et al., 2002). Particularly, the downregulation of *SLC12A1* in the gobies could be an adaptive response to restore any changes in GABA function (Grosell, 2011; Hübner & Holthoff, 2013) similar to what has been observed with the potassium-chloride co- transporter 2 (KCC2) gene in the damselfish *Acanthochromis polyacanthus* exposed to elevated CO_2_ (Schunter et al., 2018). Therefore, gobies inhabiting the naturally acidified waters of Vulcano Island possibly use alternative strategies to ensure proper functioning of neural signalling pathways reliant on maintenance of stable ionic gradients which are altered by pH buffering processes.

The circadian rhythm is a key pathway regulating nearly all physiological processes and all core transcripts in this pathway showed differential expression. CO_2_ induced changes in the circadian rhythm have been repeatedly observed across various species and are possibly linked to changes in GABA receptor activity (Lee et al., 2021). The circadian pathway enables organisms to anticipate and respond to environmental changes which is evolutionarily advantageous (Ayyar & Sukumaran, 2021; Dmitriev & Mangel, 2000; Prokkola & Nikinmaa, 2018). Changes in the expression of circadian genes in response to elevated CO_2_, have been suggested to result in an amplitude or phase shift of the circadian clock providing an adaptive advantage by increasing flexibility of physiological processes (Kang et al., 2022; Lee et al., 2021; Schunter et al., 2016, 2021). We found evidence of potential circadian regulation of metabolism in gobies in response to elevated environmental CO_2_. In addition, *NOCT,* an immediate early gene with deadenylase activity was upregulated in gobies from CO_2_ seep. *NOCT* functions as a mediator of post-transcriptional circadian regulation of metabolic processes and can also respond to changes in the external environment (Stubblefield et al., 2012). Increased expression of *NOCT* in individuals from the CO_2_ seep combined with DE of all core circadian genes suggests rhythmic regulation of energy metabolism, potentially enabling gobies to meet the increased energetic demands of living in an environment with higher than ambient levels of CO_2_. Given the ubiquity of the circadian clock, there is a need for further research to understand the biological implications of the effect of elevated CO_2_ on the circadian rhythm and its impacts on neuronal signal transduction and other downstream physiological processes.

In addition to the above-mentioned rhythmic regulation of metabolism, we found an overall increase in expression of transcripts involved in ATP production in gobies from the CO_2_ seep. Interestingly, transcripts involved in fatty-acid biosynthesis, an anabolic process was downregulated. The constant need to maintain acid-base balance and homeostasis while minimising oxidative stress in a high CO_2_ environment can be energetically expensive (Cattano et al., 2018; Kelly & Hofmann, 2013; Strader et al., 2020). Decreased expression of transcripts involved in fatty acid synthesis could be an alternative strategy of gobies living in the seep to facilitate allocation of acetyl-CoA, a key metabolite that intersects with glycolysis, TCA cycle, fatty acid beta-oxidation and fatty acid synthesis (Currais et al., 2019), for use in energy producing metabolic pathways. This in addition to up-regulation of *SLC16A1* and *SLC25A22*, which are also involved in transport of key metabolic intermediates, suggests that gobies selectively regulate metabolic pathways to ensure increased energy production. Additionally, all transcripts encoding hexose/glucose transporters, with the exception of *SLC45A1* were also upregulated in gobies from the CO_2_ seep which could facilitate increased availability of glucose in the brain. This is particularly important as glucose is the primary substrate for energy production in the brain. Despite being a glucose transporter, *SLC45A1* was selectively downregulated probably because it is a sugar-H^+^ symporter (Shimokawa et al., 2002) and hence facilitates H^+^ import into neuronal cells thereby increasing intracellular pH which could be detrimental. Therefore, gobies have potentially acclimated to a high CO_2_ environment by selective regulation of genes involved in metabolism to increased energy production. Changes in the energy budgets of fish living in acidified waters could result in physiological trade-offs (Kelly & Hofmann, 2013). However, the increased metabolic capacity of gobies observed here combined with CO_2_ driven habitat shifts resulting in enrichment of key trophic resources at the CO_2_ seep (Nagelkerken et al., 2015) potentially enables them to meet increased energy requirements. This aids in maintaining pH homeostasis and mitigating any potential cellular damage due to elevated CO_2_ levels without a significant trade-off in the energy available for other biological processes. In fact, the CO_2_ vents have a higher abundance of gobies compared to nearby control sites (Mirasole et al., 2019; Nagelkerken et al., 2015; Spatafora et al., 2022). At other CO_2_ seeps at White Island, New Zealand, a similar habitat shift with net resource enrichment can be found and the common triplefin *F. lapillum* also exhibit a higher abundance with a similar increase in metabolic capacity (Nagelkerken et al., 2015; Petit-Marty et al., 2021). Therefore, gobies similar to the common triplefin, might benefit from the habitat shifts at CO_2_ seeps and show an adaptive response to prolonged CO_2_ exposure with an overall increase in metabolism.

Our findings show molecular signatures of acclimation in the anemone goby to the acidified waters of Vulcano Island. The gobies effectively regulate disturbances in acid-base balance, maintain cellular homeostasis and mitigate OA induced oxidative stress. Gobies also showed potential adaptive mechanisms to maintain proper functioning neurological signalling pathways, which could be disrupted as a consequence of altered ionic composition due to pH buffering processes. Although these processes can be energetically expensive potentially resulting in trade-offs, the overall increase in metabolism seen in gobies from the CO_2_ seep suggests that they meet the increased energy demands associated with living in an environment with higher than ambient CO_2_ concentrations. This could be further facilitated by net trophic resource enrichment at the CO_2_ seep. The increased population density of gobies at the CO_2_ seep in Vulcano Island reported in previous studies combined with evidence of acclimation to elevated CO_2_ environments (potentially mediated by developmental plasticity) found in our study indicates that gobies can effectively mitigate the behavioural effects of elevated CO_2_ exposure. Given that a wide range of species show developmental plasticity (Munday et al., 2013) this could be a vital process in decreasing species vulnerability and result in establishment of sufficient standing phenotypic variation within populations which could become targets for selection to act upon in the future as OA continues to occur at a whole ocean scale.

## Author contributions

AM conceived the project with input from CS and TR. AM conducted the fieldwork and dissected the fish. SV provided the funding for fieldwork. SS and CS carried out the RNA extractions, designed the RNA-Seq analysis pipeline and drafted the manuscript. All authors contributed to data interpretation and editing the manuscript. All authors read and approved the manuscript.

## Acknowledgments

We thank Dr. Juan Diego Gaitan-Espitia and Dr. Kang Jingliang for providing guidance and feedback on the de novo transcriptome assembly. Ethics

All sample collection were carried out in accordance with the institutional and national law guidelines concerning the use of animals in research.

## Data accessibility

RNA-seq raw sequences and the *de novo* assembled transcriptome assemblies have been deposited in NCBI under BioProject PRJNA869880.

