## Supplementary Figures for "Brain transcriptome of gobies inhabiting natural CO2 seeps reveal acclimation strategies to long-term acidification"

**Supplemenary Figure 1:**

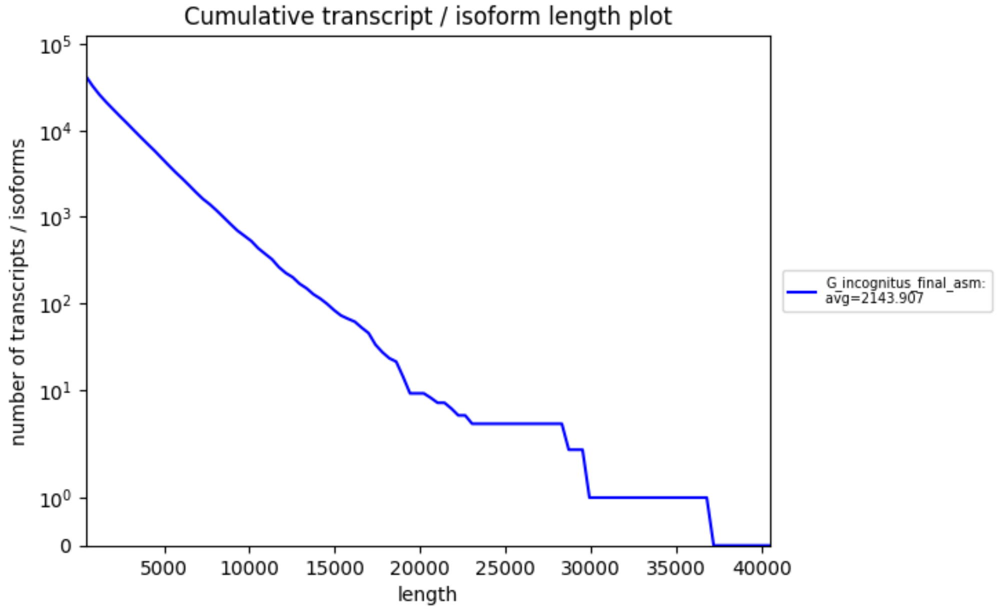


**Figure S1:** Cumulative length distribution of all assembled transcripts in the *de novo* transcriptome. The *de novo* assembled transcriptome consisting of 43,349 contigs (transcripts) has an average length of 2,143.9 bp with an N50 value of 3,545, L50 value of 7,998 and 61% of the contigs over 1Kbp in length.

**Supplemenary Figure 2:**

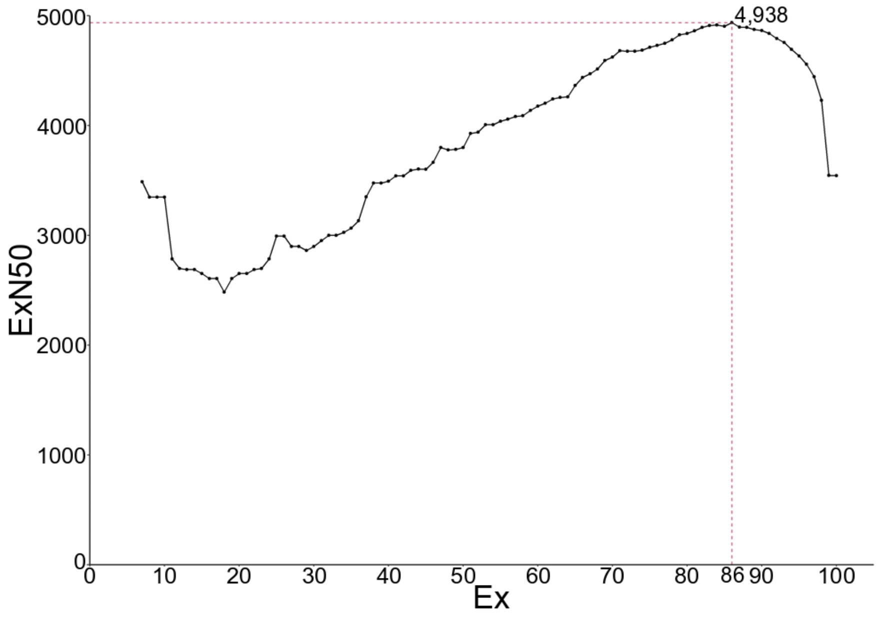

**Figure S2:** Expression-informed N50 (ExN50) calculated across different expression percentiles (Ex) using the TMM normalised counts. The highest ExN50 value (4,938 bp) was obtained when including transcripts representing 86% of total normalised expression indicating good coverage of longer transcripts in the *de novo* assembly.

**Supplemenary Figure 3:**

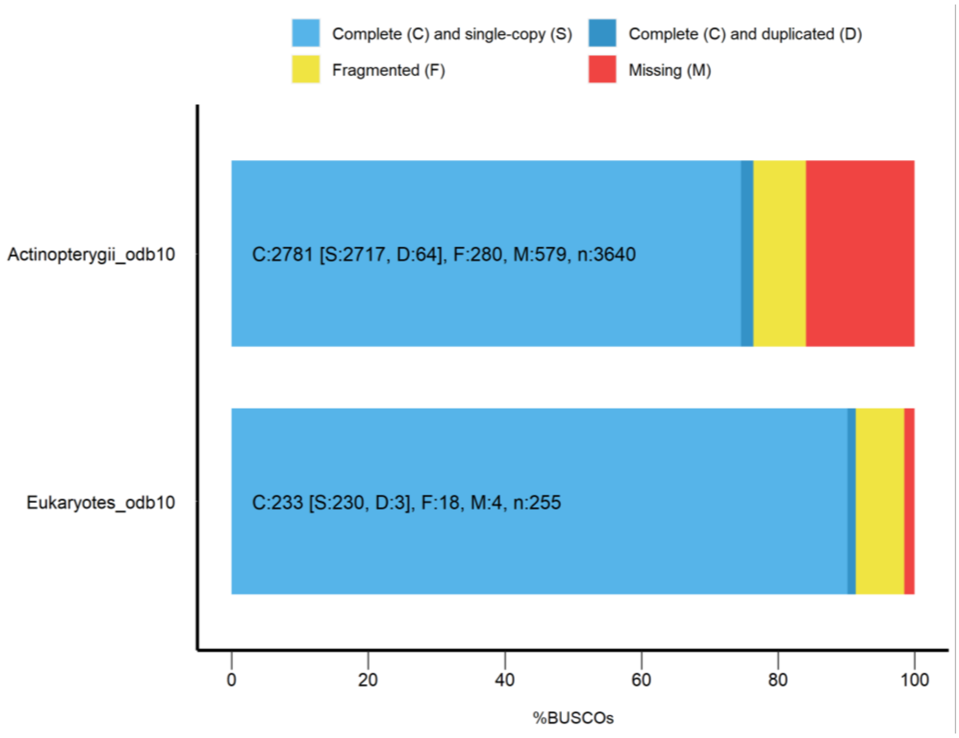


**Figure S3:** BUSCO assessment of assembly completeness using the Actinopterygii_odb10 and Eukaryota_odb10 core gene dataset. The analyses revealed that the *de novo* assembled transcriptome had a high gene completeness with 84% and 98% of genes from the Actinopterygii and Eukaryota library recovered respectively

**Supplemenary Figure 4:**

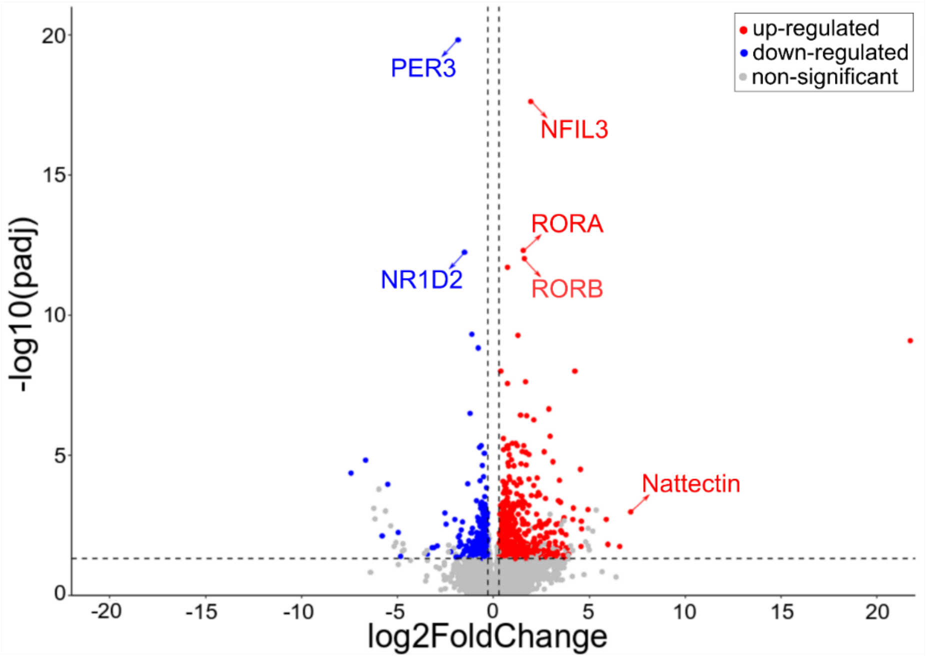


**Figure S4:** Volcano plot showing log2 fold change in transcript expression levels between individuals from control and $CO_2$ seep (LPH) sites on the x-axis (positive values are up-regulated and negative values are down-regulated in fish from the $CO_2$ seeps sites) and the negative log10 of FDR corrected p-values on the y-axis. Transcripts that are not significantly DE are in grey, transcripts that are significantly DE are in red (up-regulated) and blue (downregulated).

**Supplemenary Figure 5:**

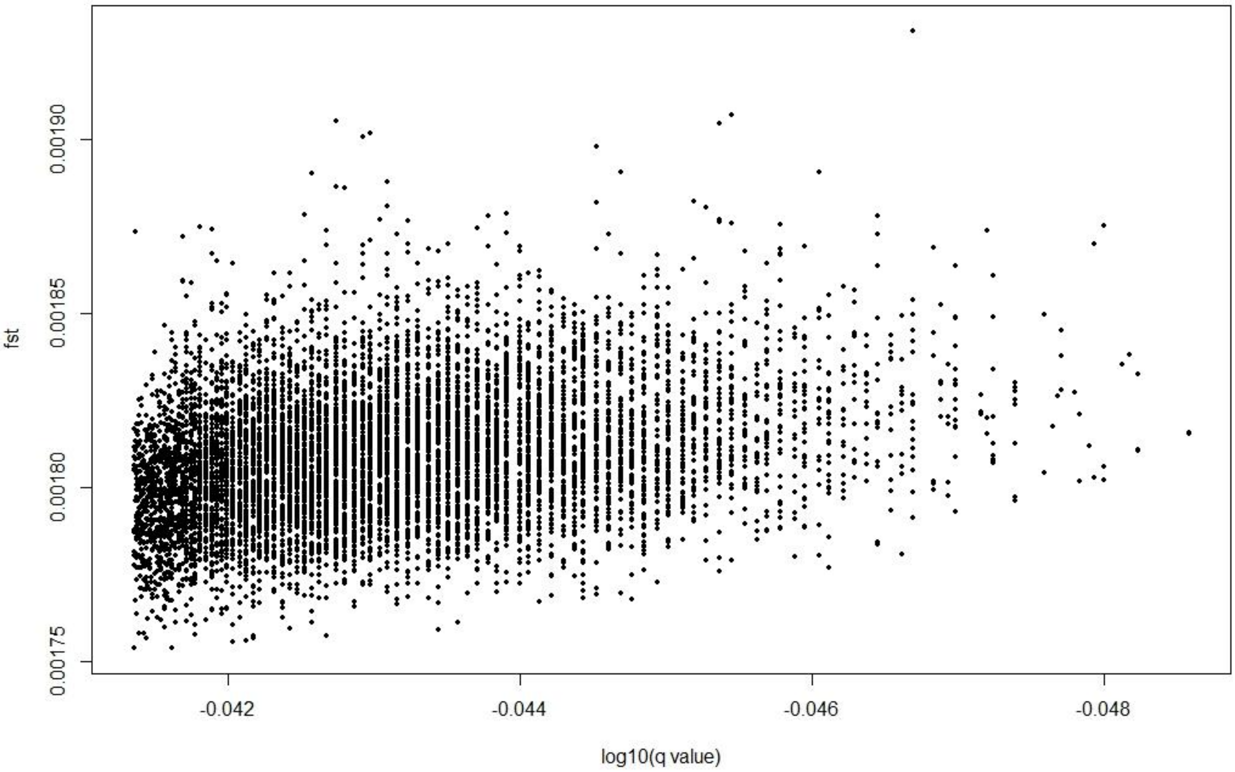


**Figure S5:** Detection of putative outlier loci between the samples from the $CO_2$ seep and control sites using the Bayesian based BayeScan program. The graph represents the $F_{ST}$ values against the corresponding log10(FDR corrected p-value(q value)) for each loci. There were no significant outlier loci detected between the samples from the $CO_2$ seep and control sites.
